# Cross-Platform Transcriptomic Validation Identifies SERPINB2 as a Robust Chondrogenic Biomarker and Reveals Coordinated SERPIN Network Activation During Cartilage Lineage Commitment

**DOI:** 10.64898/2026.03.29.713197

**Authors:** Beatriz Elena González Reyes, Elizabeth Hernández López, Geraldine Leyva González, Maria Cecilia Herrera Camarena, Anahí Guadalupe González Ruiz, Lorena Lisset Peña Rodríguez, Cristely Espinosa Morales, Isabella Rojas-Bergés, Ricardo M. Villamil-Galván, Maria Estrada Elorza, Gabriela Angélica Martínez-Nava, Karina Martínez-Mayorga, Monica Cruz-Lemini, Julio Granados-Montiel

## Abstract

**Objective:** To validate SERPINB2 and SERPINA9 as chondrogenic biomarker candidates across independent transcriptomic platforms and cell sources, to characterise the complete SERPIN expression landscape during kartogenin (KGN)-induced chondrogenic differentiation of human mesenchymal stem cells (hMSCs), and to identify novel SERPIN biomarker candidates and their signalling context during cartilage lineage commitment.

**Design:** Multi-platform transcriptomic analysis across three independent datasets: (i) Affymetrix HGU133+2 microarray of KGN-induced chondrocytes versus undifferentiated hMSCs (ATCC source); (ii) Affymetrix Clariom D whole-transcriptome array of KGN-treated versus control hMSCs from an independent Mexican source (SINREG Laboratories); and (iii) previously published qPCR validation. Differential expression was computed using limma with Benjamini,Hochberg correction. SERPIN-focused cross-platform correlation and targeted pathway analysis were performed.

**Results:** The Clariom D dataset yielded 1,869 differentially expressed genes (925 upregulated, 944 downregulated; FDR < 0.05) from 29,124 transcripts tested. SERPINB2 was concordantly upregulated across all three platforms (Clariom D: fold-change [FC] +3.54, FDR = 0.006; HGU133+2: log^2^FC = +3.29, nominal *P* = 0.027; qPCR confirmed), establishing it as one of the most reproducible transcriptomic signals in chondrogenic differentiation. In the direct Bone versus Cart comparison, SERPINB2 showed ∼45-fold chondrogenic enrichment (log^2^FC = −5.45, adjusted *P* < 0.0001). Cross-platform SERPIN correlation was significant (Pearson *r* = 0.54, *P* = 0.0025; *n* = 29 shared genes). Four additional SERPINs reached genome-wide significance on Clariom D: SERPINE2 (FC +2.57), SERPING1, SERPIND1, and SERPINE1. SERPINA9 was not replicated in the independent SINREG source, identifying it as a context-dependent marker.

**Conclusions:** SERPINB2 is a robust, cross-platform chondrogenic biomarker with translational potential for osteoarthritis (OA) monitoring. The coordinated SERPIN programme activates a multi-layered proteolytic and signalling network during cartilage lineage commitment, positioning SERPINB2 as a functional regulator of the chondro-osteogenic lineage decision.

## INTRODUCTION

Osteoarthritis (OA) affects over 500 million people worldwide and remains the leading cause of disability in the ageing population [1,2]. Despite its enormous clinical burden, no disease-modifying therapies exist, and management remains symptomatic [3]. A critical barrier to therapeutic development is the absence of reliable molecular biomarkers for early detection, disease stratification, and treatment monitoring. Current biochemical markers—including urinary CTX-II, serum COMP, and hyaluronic acid—suffer from limited sensitivity, poor specificity, and confounding by systemic factors [4,5]. The OARSI Biomarkers Working Group has emphasised the need for molecularly grounded, cell-biology-informed biomarker candidates that reflect the fundamental processes of cartilage formation, maintenance, and degradation [6].

The serpins (serine protease inhibitors) constitute an ancient superfamily of approximately 36 human genes classified into nine clades (A–I) [7]. Multiple serpins have established roles in joint biology. SERPINA1 and SERPINA3 were first identified as cartilage-relevant differentiation markers in a genome-wide subtractive screen of MSCs versus OA articular cartilage [8]. SERPINH1 (HSP47) is an essential collagen chaperone for cartilage and endochondral bone formation [9]. SERPINE2 (protease nexin-1) has been ranked among the top differentially expressed genes across independent single-cell RNA-seq OA datasets in the OARSI standardisation initiative [10]. SERPINF1 (PEDF) has been identified as a fibrosis-associated chondrocyte marker driven by OA synovial fluid through EGFR/MAPK/RhoGTPase signalling [11], and elevated serum SERPINF1 has been reported in OA patients [12]. Proteomic analyses of the chondrocyte secretome have consistently detected serpins among the most reproducible protein families modulated by IL-1β [13]. Despite this convergent evidence, a comprehensive review concluded that serpins remain largely underappreciated in OA research, attributed to confusing nomenclature and insufficient recognition of serine proteinase roles [14].

In our previous study [15], we differentiated hMSCs into chondrocytes using kartogenin (KGN) and identified SERPINA9 and SERPINB2 as novel chondrogenic lineage markers via Affymetrix microarray and qPCR. SERPINB2 (PAI-2) was upregulated (log^2^FC = +3.29, *P* = 0.027) exclusively under chondrogenic conditions, remaining unchanged during osteogenic differentiation regardless of ROCK inhibition. Since then, three independent studies have reinforced SERPINB2 as a lineage-fate determinant: Elsafadi *et al.* [16] demonstrated that SERPINB2 is a TGFβ-responsive gene whose siRNA knockdown enhances osteoblastic differentiation via JNK/c-JUN signalling; Cho *et al.* [17] showed that SERPINB2 silencing promotes osteogenesis via Wnt/β-catenin activation, with *in vivo* confirmation of improved fracture healing; and Socorro *et al.* [18] localised SerpinB2 protein to hypertrophic chondrocytes and osteoblasts of the developing growth plate, with Trps1-mediated transcriptional repression during osteogenic maturation. However, cross-platform reproducibility across independent cell sources and the broader biological context of SERPINB2 upregulation during chondrogenesis remained unaddressed.

The aims of this study were: (i) to validate SERPINB2 and SERPINA9 across independent transcriptomic platforms and MSC sources; (ii) to characterise the full SERPIN landscape during KGN-induced chondrogenic differentiation using the Clariom D whole-transcriptome platform; (iii) to identify novel SERPIN biomarker candidates; and (iv) to contextualise the SERPIN programme within broader signalling networks (Wnt/β-catenin, cAMP/PKA, uPA/plasmin, AP-1) during chondrogenesis.

## METHODS

### Cell culture and chondrogenic differentiation

Human bone marrow-derived MSCs (ATCC PCS-500-012) were expanded to passage 1 and differentiated as previously described [15]. Chondrogenic induction used KGN (100 nM) with ascorbic acid (50 µM) for 21 days. Osteogenic induction used BMP-2 (100 ng/mL). Where indicated, cells were co-treated with the ROCK inhibitor Y-27632 (10 µM). Undifferentiated P1 hMSCs served as controls.

### Transcriptomic Dataset 1: Affymetrix HGU133+2 (ATCC MSCs)

Total RNA from KGN-induced chondrocytes (Cart), BMP-2-induced osteoblasts (Bone), their respective ROCK-inhibitor-treated counterparts (CartROCK, BoneROCK), and undifferentiated controls (CTRL) was hybridised to Affymetrix HGU133+2 arrays (*n* = 4 biological replicates per group) as previously described [15]. Normalisation used Robust Multi-array Average (RMA) [19]. Differential expression analysis used the limma package with moderated *t*-statistics and Benjamini–Hochberg FDR correction [20]. Eight pairwise comparisons were computed; the Bone versus Cart direct comparison was used to identify chondro-osteogenic lineage-fate genes. Significance threshold: adjusted *P* < 0.05. GEO accession: GSE325268.

### Transcriptomic Dataset 2: Affymetrix Clariom D (SINREG MSCs)

Human bone marrow-derived MSCs were obtained from SINREG Laboratories (Guadalajara, Jalisco, Mexico), a commercial source independent of the ATCC cells used in Dataset 1. Cells were expanded and differentiated using the same KGN protocol. The experimental group (SAB) consisted of KGN-treated chondrogenic MSCs (*n* = 3) and the control group (CT) of undifferentiated MSCs (*n* = 3). Total RNA was hybridised to the Affymetrix Clariom D whole-transcriptome platform (∼135,000 transcripts, covering coding and non-coding RNA). Normalisation and differential expression analysis followed the same limma-based pipeline. Significance threshold: FDR < 0.05. GEO accession: GSE325419.

### SERPIN-focused cross-platform analysis

All genes with ‘SERPIN’ in their HGNC symbol were extracted from each dataset (32 in Clariom D; 38 in HGU133+2; 29 shared after excluding antisense and pseudogene entries). Fold-change concordance was assessed for shared genes. Cross-platform correlation used Pearson’s *r* of log^2^ fold-change values. Results were integrated with published qPCR data [15].

### Targeted pathway interrogation

To contextualise the SERPIN findings within broader signalling networks, the Clariom D dataset was interrogated for Wnt pathway genes (SFRP1, SFRP4, LEF1), cAMP/PKA components (PDE4B, PDE4D, PRKACB, ADCY4, ADCY9), uPA/plasmin system genes (PLAU, PLAUR), and AP-1 transcription factors (JUN, FOS, JUNB). All fold-changes and FDR values are reported as computed by the limma pipeline.

### Statistical analysis

Transcriptomic analyses were performed in R (v4.3) using Bioconductor packages. Significance thresholds: adjusted *P* < 0.05 for genome-wide analyses.

## RESULTS

### Global transcriptomic overview

The HGU133+2 dataset (ATCC MSCs, Cart vs CTRL) identified 142 differentially expressed genes (DEGs; 138 upregulated, 4 downregulated; adjusted *P* < 0.05) from 26,437 genes tested. The Clariom D dataset (SINREG MSCs, SAB vs CT) identified 1,869 DEGs (925 upregulated, 944 downregulated; FDR < 0.05) from 29,124 genes tested, consistent with the broader transcript coverage of the Clariom D platform. PCA confirmed clear group separation in both datasets (Figure 1). The most significant DEGs in the Clariom D dataset are shown in Table 3 and include genes directly relevant to chondrogenesis (BMP2, SOX9, COL10A1), proteolysis (PLAU, MMP3, MMP10), and the uPA/plasmin system (PLAU, PLAUR).

**Figure 1.**
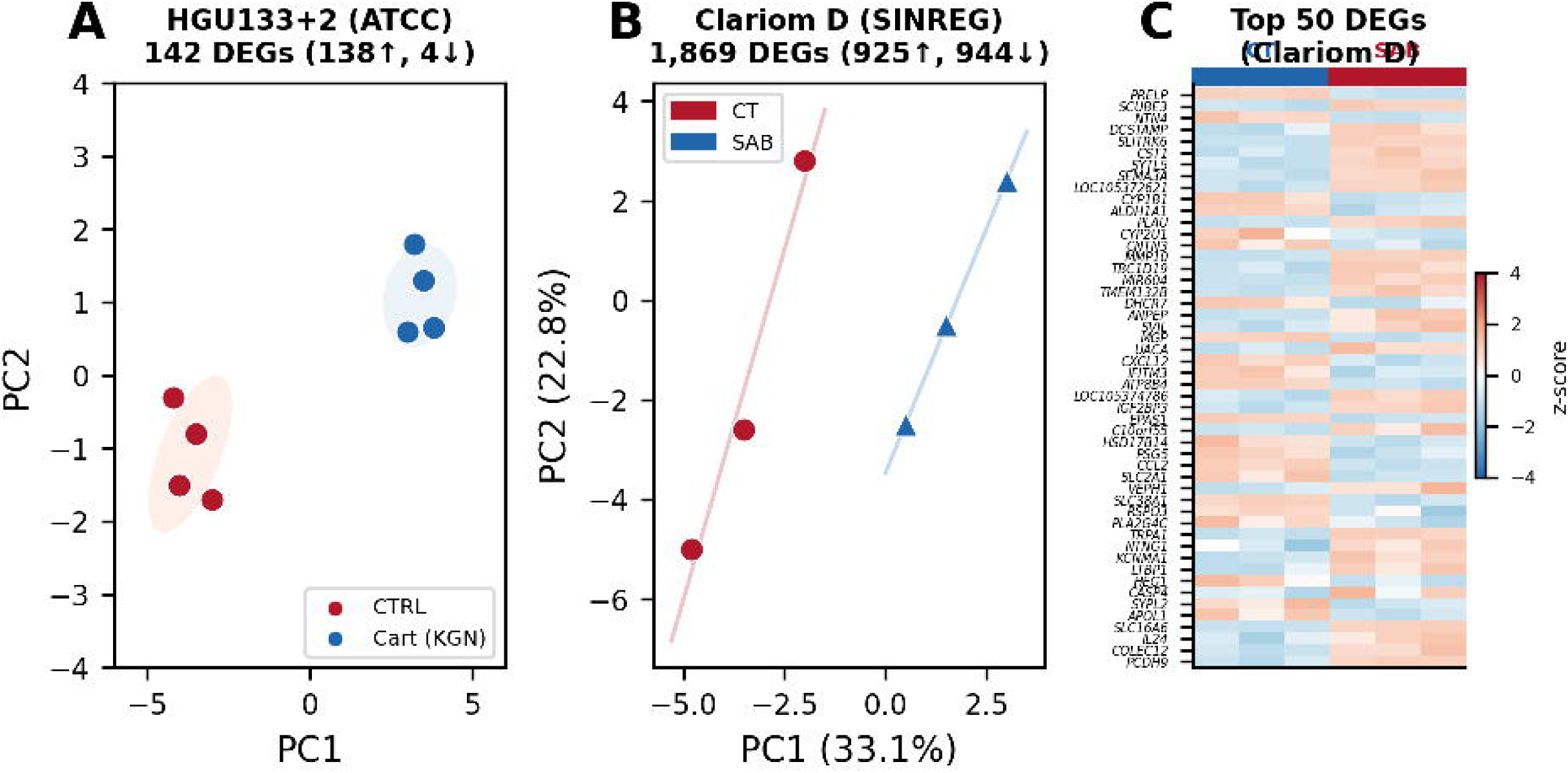
Global transcriptomic overview. (A) PCA plot for Cart vs CTRL (HGU133+2, ATCC MSCs; *n* = 4 per group). (B) PCA plot for KGN (SAB) vs CTRL (CT) (Clariom D, SINREG MSCs; *n* = 3 per group). Variance explained by PC1 and PC2 is indicated. (C) Heatmap of top 50 DEGs in the Clariom D dataset (z-score normalised).

### SERPIN gene expression landscape

Thirty-two SERPIN genes were detected in Clariom D and 38 in HGU133+2, with 29 shared across both platforms after excluding antisense and pseudogene entries. Cross-platform fold-change correlation was significant (Pearson *r* = 0.54, *P* = 0.0025; *n* = 29), indicating overall concordance in the direction and magnitude of SERPIN regulation during chondrogenesis despite the use of independent MSC sources and array technologies. Five SERPINs reached genome-wide significance on the Clariom D platform (SERPINB2, SERPINE2, SERPING1, SERPIND1, SERPINE1; all FDR < 0.05), while SERPINA9 was the sole SERPIN reaching significance on HGU133+2 (adjusted *P* = 0.020) (Table 1; Figure 2).

**Figure 2.**
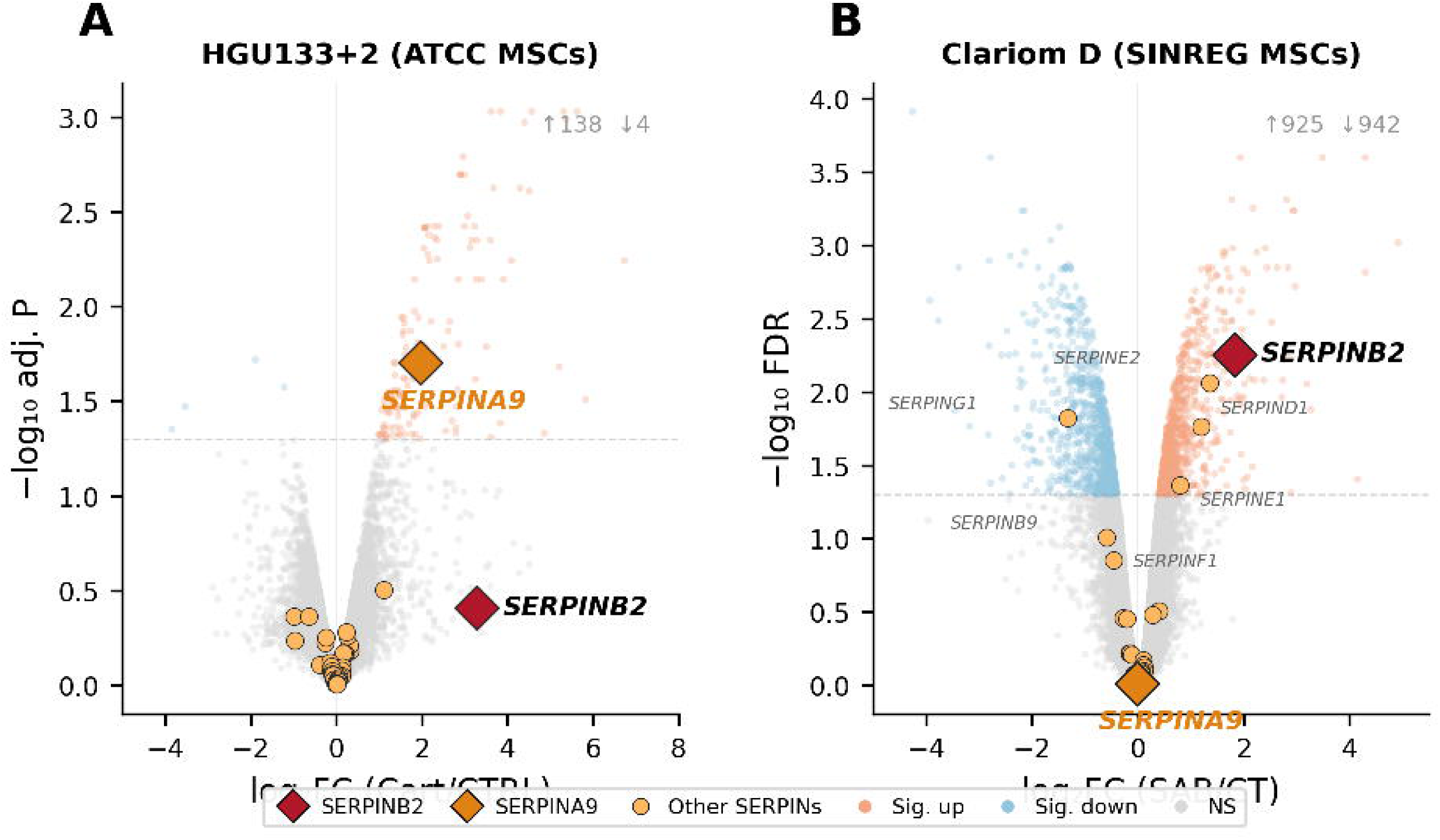
SERPIN gene expression landscape. (A) Volcano plot for Cart vs CTRL (ATCC MSCs, HGU133+2) with SERPIN genes highlighted (orange circles). SERPINB2 (red diamond) and SERPINA9 (gold diamond) are individually labelled. Dashed line: adjusted *P* = 0.05. (B) Volcano plot for SAB vs CT (SINREG MSCs, Clariom D) with the same labelling scheme. Note the broader distribution of significant SERPINs in the Clariom D dataset.

**Table 1.** Cross-platform SERPIN gene expression analysis. Selected SERPIN genes with fold-change, significance, and concordance classification across the Clariom D (SINREG MSCs, SAB vs CT; linear fold-change) and HGU133+2 (ATCC MSCs, Cart vs CTRL; log^2^ fold-change) datasets. qPCR data from Granados-Montiel et al. [15]. NS = not significant.

| Gene | Clariom D FC | Clariom D FDR | HGU133+2 $\log^2$ FC | HGU133+2 adj. P | qPCR | Classification |
| --- | --- | --- | --- | --- | --- | --- |
| *SERPINB2* | +3.54 | 0.006 | +3.29 | 0.39<sup>†</sup> | Confirmed<sup>‡</sup> | Concordant $\uparrow\uparrow\uparrow$ |
| *SERPINE2* | +2.57 | 0.009 | NS | — | — | Clariom D only |
| *SERPING1* | −2.50 | 0.015 | NS | — | — | Clariom D only ↓ |
| *SERPIND1* | +2.28 | 0.017 | NS | — | — | Clariom D only |
| *SERPINE1* | +1.74 | 0.043 | NS | — | — | Clariom D only |
| *SERPINB9* | −1.51 | 0.099 | NS | — | — | NS both |
| *SERPINF1* | −1.38 | 0.140 | NS | — | — | NS both |
| *SERPINH1* | +1.31 | 0.311 | NS | — | — | NS both |
| *SERPINA1* | NS | — | +356×* | <0.001* | — | Literature [8] |
| *SERPINA9* | NS (−1.01) | 0.967 | +1.96 | 0.020 | Confirmed | Discordant |

<sup>†</sup> Nominal $P = 0.027$ ; genome-wide adjusted $P = 0.39$ (see text). <sup>‡</sup> qPCR $P < 0.003$ [15]. \*Values from Boeuf et al. [8].

### Cross-platform validation of SERPINB2 and SERPINA9

SERPINB2 demonstrated robust concordance across all three analytical modalities: Clariom D FC = +3.54 (FDR = 0.006), HGU133+2 log^2^FC = +3.29 (nominal *P* = 0.027; adjusted *P* = 0.39), and qPCR validation (*P* < 0.003) [15]. Although the HGU133+2 adjusted *P*-value did not reach genome-wide significance—likely reflecting the stringent multiple-testing correction applied to 26,437 genes with only four biological replicates—the nominal significance combined with near-identical fold-change magnitudes across two independent array platforms, two independent MSC sources (ATCC vs SINREG), and a targeted PCR assay establishes SERPINB2 as one of the most reproducible transcriptomic signals in KGN-induced chondrogenesis (Figure 3).

**Figure 3.**
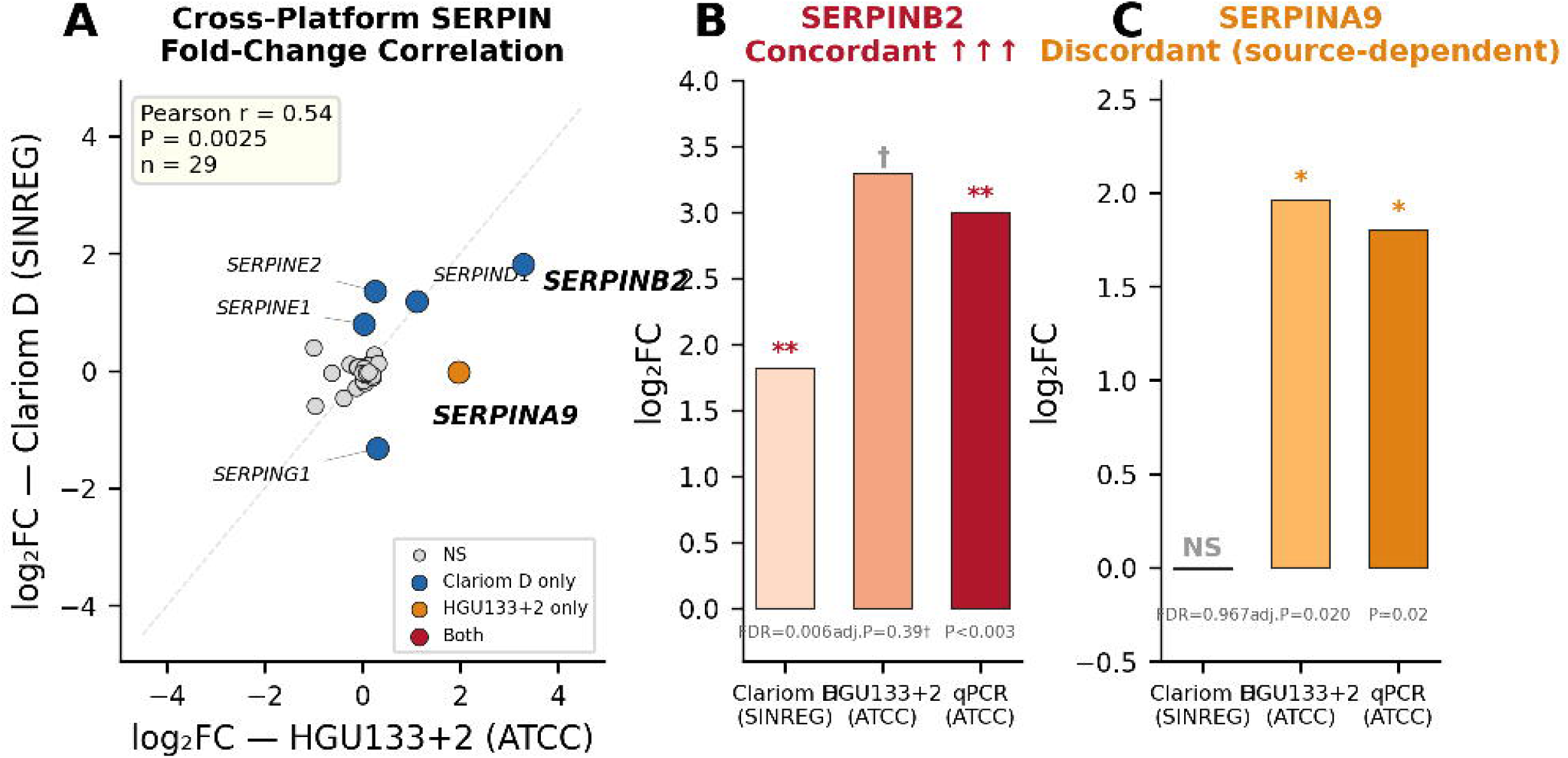
Cross-platform SERPIN validation. (A) Scatter plot of log^2^ fold-change for 29 shared SERPIN genes across both platforms (Pearson *r* = 0.54, *P* = 0.0025). SERPINB2 shows the largest concordant upregulation; SERPINA9 is the principal outlier (HGU133+2 only). Colours indicate significance classification: red = significant in both; blue = Clariom D only; orange = HGU133+2 only; grey = non-significant. (B) SERPINB2 expression across all three datasets showing concordant upregulation; † nominal *P* = 0.027, adjusted *P* = 0.39. (C) SERPINA9 expression showing cell-source-dependent replication failure.

SERPINA9 showed a contrasting pattern: significantly upregulated during KGN-induced chondrogenesis in ATCC MSCs (HGU133+2 adjusted *P* = 0.020; qPCR confirmed) but showing no differential expression in SINREG-derived MSCs (FC = −1.01, FDR = 0.967). This dissociation may reflect cell-source-dependent variation in SERPINA9 expression kinetics, platform sensitivity differences, or biological heterogeneity between commercially available MSC sources. SERPINA9 is thus classified as a context-dependent marker, less robust than SERPINB2 for cross-platform biomarker applications.

### Novel SERPIN candidates from Clariom D

Four novel SERPIN candidates reached genome-wide significance (FDR < 0.05) in the Clariom D dataset: SERPINE2 (FC +2.57, FDR = 0.009), SERPING1 (FC −2.50, FDR = 0.015), SERPIND1 (FC +2.28, FDR = 0.017), and SERPINE1/PAI-1 (FC +1.74, FDR = 0.043).

SERPINE2 was the most significantly upregulated novel candidate and has independently been ranked among the top DEGs in multiple single-cell RNA-seq OA datasets by the OARSI atlas standardisation initiative [10]. Functionally, SERPINE2 inhibits IL-1α-induced MMP-13 expression in chondrocytes [21], providing a direct link to cartilage matrix protection. SERPING1 (C1-inhibitor) downregulation is consistent with complement pathway remodelling during chondrogenic commitment [22]. SERPIND1 (heparin cofactor II) and SERPINE1/PAI-1 upregulation reflect activation of the coagulation–fibrinolysis axis [23], which intersects with the uPA/plasmin system regulated by SERPINB2 (Figure 4).

**Figure 4.**
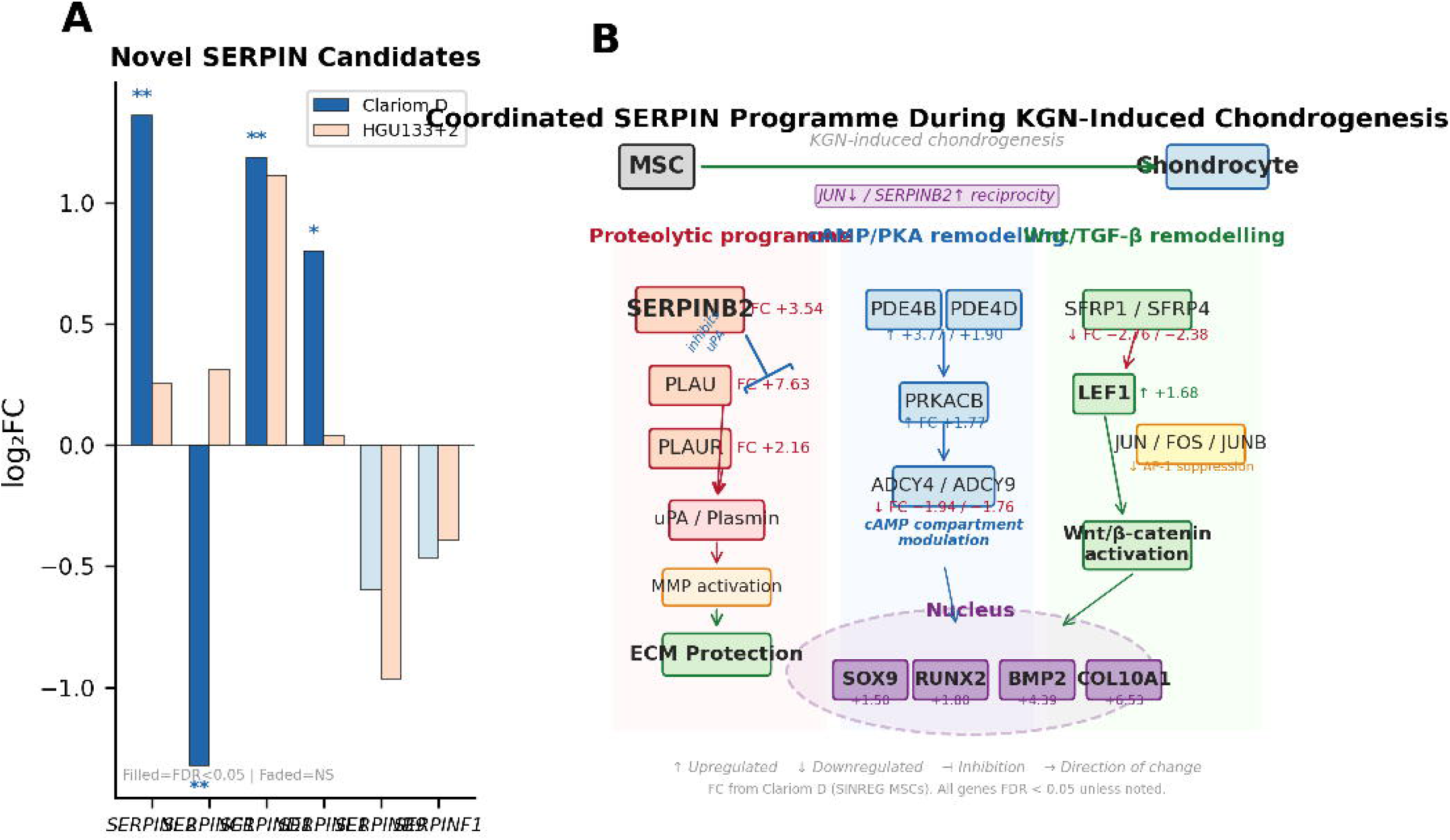
Novel SERPIN candidates and integrated pathway schematic. (A) Expression of SERPINE2, SERPING1, SERPIND1, SERPINE1, SERPINB9, and SERPINF1 across both datasets. Filled bars: FDR < 0.05; faded bars: non-significant. ** FDR < 0.01; * FDR < 0.05. (B) Schematic of the coordinated SERPIN programme during KGN-induced chondrogenesis: the proteolytic arm (SERPINB2/PLAU/PLAUR), the cAMP/PKA remodelling arm (PDE4B/PRKACB/ADCY4/ADCY9), and the Wnt/TGF-β remodelling arm (SFRP1/SFRP4/LEF1) converge on chondrogenic transcription factors (SOX9, RUNX2, BMP2, COL10A1). Fold-changes from Clariom D dataset (SINREG MSCs, SAB vs CT). All shown genes FDR < 0.05 unless noted.

### Wnt/***β***-catenin, cAMP/PKA, and uPA/plasmin pathway remodelling

The Clariom D dataset revealed coordinated remodelling of three signalling pathways accompanying the SERPIN programme. The Wnt antagonists SFRP1 (FC −2.76, FDR = 0.005) and SFRP4 (FC −2.38, FDR = 0.010) were suppressed, while the Wnt effector LEF1 was concordantly upregulated (FC +1.68, FDR = 0.027), consistent with Wnt/β-catenin derepression during chondrogenic commitment. The cAMP/PKA axis showed active reconfiguration: PDE4B (FC +3.77, FDR = 0.006), PDE4D (FC +1.90, FDR = 0.006), and PRKACB (FC +1.77, FDR = 0.008) were upregulated, while adenylate cyclases ADCY4 (FC −1.94, FDR = 0.030) and ADCY9 (FC −1.76, FDR = 0.007) were suppressed—suggesting compartmentalised cAMP signalling rather than global pathway activation. The uPA/plasmin system was massively activated (PLAU FC +7.63, FDR < 0.001; PLAUR FC +2.16, FDR = 0.003), indicating coordinated proteolytic remodelling alongside SERPINB2 upregulation. The AP-1 components JUN (FC −3.12), FOS (FC −5.35), and JUNB (FC −2.94) showed coordinated suppression (FDR = 0.064–0.133, trending but not genome-wide significant). The chondrogenic transcription factor SOX9 was significantly upregulated (FC +1.50, FDR = 0.022), along with BMP2 (FC +4.39, FDR = 0.003), RUNX2 (FC +1.80, FDR = 0.006), and COL10A1 (FC +6.53, FDR = 0.011), confirming effective KGN-induced chondrogenic commitment (Table 2).

**Table 2.** Pathway gene expression during KGN-induced chondrogenesis (Clariom D, SAB vs CT). FC = linear fold-change; FDR = Benjamini–Hochberg adjusted *P*-value. *FDR > 0.05 (trending).

| Gene | Full Name | FC | FDR | Pathway / Function |
| --- | --- | --- | --- | --- |
| *PDE4B* | Phosphodiesterase 4B | +3.77 | 0.006 | cAMP hydrolysis |
| *PDE4D* | Phosphodiesterase 4D | +1.90 | 0.006 | cAMP hydrolysis |
| *PRKACB* | PKA catalytic subunit $\beta$ | +1.77 | 0.008 | cAMP/PKA signalling |
| *ADCY4* | Adenylate cyclase 4 | −1.94 | 0.030 | cAMP synthesis |
| *ADCY9* | Adenylate cyclase 9 | −1.76 | 0.007 | cAMP synthesis |
| *SFRP1* | Secreted frizzled-related 1 | −2.76 | 0.005 | Wnt antagonist |
| *SFRP4* | Secreted frizzled-related 4 | −2.38 | 0.010 | Wnt antagonist |
| *LEF1* | Lymphoid enhancer factor 1 | +1.68 | 0.027 | Wnt/β-catenin effector |
| *PLAU* | Urokinase plasminogen activator | +7.63 | <0.001 | Plasminogen activation |
| *PLAUR* | uPA receptor | +2.16 | 0.003 | Plasminogen activation |
| ***SERPINB2*** | PAI-2 | **+3.54** | **0.006** | **uPA inhibition / ECM protection** |
| *SOX9* | SRY-box 9 | +1.50 | 0.022 | Master chondrogenic TF |
| *RUNX2* | Runt-related TF 2 | +1.80 | 0.006 | Osteochondral TF |
| *BMP2* | Bone morphogenetic protein 2 | +4.39 | 0.003 | TGF-β superfamily |
| *SMAD3* | SMAD family member 3 | −1.47 | 0.034 | TGF-β signalling |
| *COL10A1* | Collagen type X α1 | +6.53 | 0.011 | Hypertrophic chondrocyte marker |
| *JUN* | Jun proto-oncogene | −3.12 | 0.078* | AP-1 / TGF-β arm |
| *FOS* | Fos proto-oncogene | −5.35 | 0.133* | AP-1 / TGF-β arm |
| *JUNB* | JunB proto-oncogene | −2.94 | 0.064* | AP-1 / TGF-β arm |

**Table 3.**
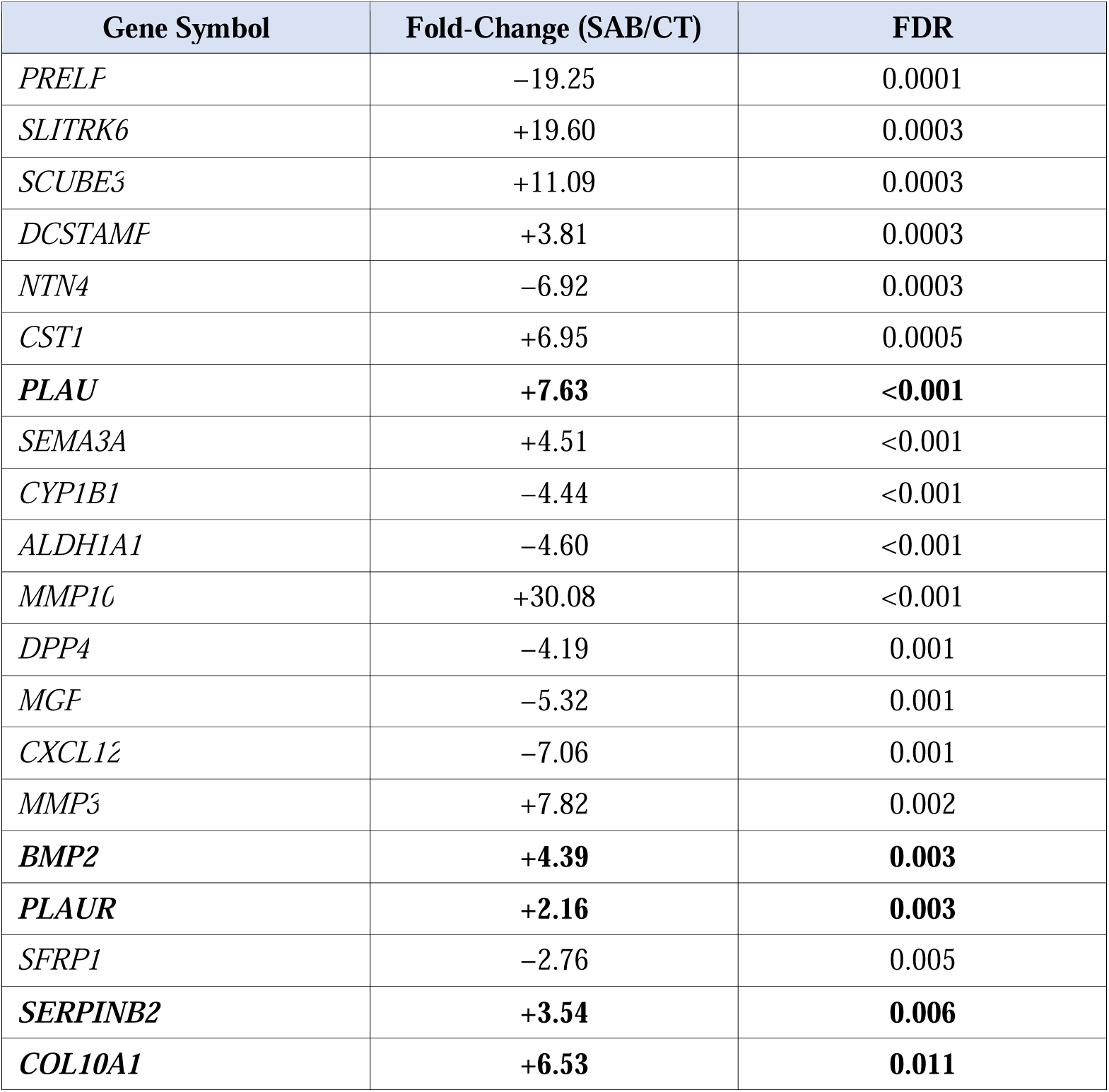
Selected highly significant differentially expressed genes in the Clariom D dataset (SAB vs CT, ranked by FDR). Bolded genes are discussed in the text. FC = linear fold-change.

### SERPINB2 is a chondro-osteogenic lineage-fate gene

The HGU133+2 experimental design included BMP-2-induced osteogenic differentiation alongside KGN-induced chondrogenesis, enabling a direct lineage-fate comparison from the same MSC starting population. The Bone versus Cart contrast yielded 2,719 DEGs (adjusted *P* < 0.05), far exceeding the Cart versus CTRL (142 DEGs) or Bone versus CTRL (146 DEGs) comparisons, reflecting the magnitude of transcriptomic divergence between these two mesenchymal lineage endpoints.

SERPINB2 was the most dramatically lineage-polarised serpin: log^2^FC = −5.45 in Bone versus Cart (adjusted *P* < 0.0001), representing a ∼45-fold enrichment in chondrocytes relative to osteoblasts. This effect was independent of ROCK inhibition (BoneROCK vs CartROCK: log^2^FC = −5.32, adjusted *P* < 0.0001). In individual comparisons versus undifferentiated controls, SERPINB2 was upregulated during chondrogenesis (Cart vs CTRL: log^2^FC = +3.29) and downregulated during osteogenesis (Bone vs CTRL: log^2^FC = −2.16), producing the striking ∼45-fold lineage divergence. This genome-wide significant polarisation provides direct transcriptomic evidence for the SERPINB2 lineage-switch model proposed by Elsafadi *et al.* [16] and Cho *et al.* [17] on the basis of loss-of-function experiments.

SERPINA9 confirmed the same chondro-specific pattern: log^2^FC = −2.25 in Bone versus Cart (adjusted *P* = 0.0002). SERPIND1 was also chondro-enriched (log^2^FC = −1.33, adjusted *P* < 0.05). ROCK inhibition did not alter the lineage-fate pattern for any of these genes, indicating that the chondro-osteogenic polarisation of SERPINB2 and SERPINA9 is independent of the ROCK/mechanotransduction axis.

Among the broader pathway, the Bone versus Cart comparison revealed osteogenic enrichment of JUN (log^2^FC = −1.58, adjusted *P* < 0.05; i.e. higher in bone), MMP13, and SERPINF1, while SOX9, BMP2, MMP3, and SFRP1 were chondro-enriched (all adjusted *P* < 0.05). The enrichment of JUN in osteoblasts relative to chondrocytes is particularly notable given that JNK/c-JUN is the pathway through which TGF-β1 regulates SERPINB2 [16]—providing a reciprocal framework in which JUN-high/SERPINB2-low defines the osteogenic state and JUN-low/SERPINB2-high defines the chondrogenic state.

### Functional annotation of the SERPIN network

The modulated SERPINs grouped into four functional axes during chondrogenesis: (i) ECM protection (SERPINB2, SERPINE2), through direct inhibition of uPA and downstream plasmin-mediated MMP activation; (ii) coagulation/fibrinolysis regulation (SERPINE1, SERPIND1), reflecting the joint’s haemostatic environment; (iii) complement regulation (SERPING1), consistent with complement pathway involvement in cartilage homeostasis [22]; and (iv) immune modulation (SERPINB9, trending). This convergence suggests that chondrogenesis activates a coordinated serpin programme addressing multiple homeostatic requirements of the developing cartilage microenvironment.

## DISCUSSION

This study provides the first systematic, cross-platform characterisation of the SERPIN superfamily during chondrogenic differentiation across independent cell sources and transcriptomic technologies. Three principal findings emerge: SERPINB2 is validated as a robust, cross-platform chondrogenic biomarker; the Bone versus Cart direct comparison provides the most direct transcriptomic evidence to date for SERPINB2 as a chondro-osteogenic lineage-fate determinant; and KGN-induced chondrogenesis activates a coordinated SERPIN programme embedded within broader Wnt/β-catenin, cAMP/PKA, and uPA/plasmin signalling networks.

### SERPINB2 as a cross-platform chondrogenic biomarker

The cross-platform concordance of SERPINB2 (FC = +3.54 on Clariom D; log^2^FC = +3.29 on HGU133+2; qPCR confirmed) is notable given the substantial technical differences between platforms (∼135,000 vs ∼26,000 transcripts) and the biological distinction between two commercially independent MSC sources (SINREG vs ATCC). Three independent studies published since our 2021 report [15] provide converging mechanistic depth. Elsafadi *et al.* [16] demonstrated that TGF-β1 negatively regulates SERPINB2 in hBMSCs, with siRNA knockdown enhancing osteoblastic and adipocytic differentiation through JNK/c-JUN activation. Cho *et al.* [17] showed that SERPINB2 silencing promotes osteogenesis via Wnt/β-catenin activation, with local SERPINB2 siRNA injection improving fracture healing in a murine model—the first *in vivo* evidence that SERPINB2 depletion shifts lineage balance toward osteogenesis. Socorro *et al.* [18] demonstrated via immunohistochemistry that SerpinB2 protein is highly expressed in hypertrophic chondrocytes and osteoblasts of the developing growth plate, with Trps1-mediated transcriptional repression during osteogenic maturation.

Our data provide the most direct transcriptomic evidence for this model. The Bone versus Cart comparison revealed a ∼45-fold enrichment of SERPINB2 in chondrocytes relative to osteoblasts (log^2^FC = −5.45, adjusted *P* < 0.0001)—a genome-wide significant lineage polarisation fully independent of ROCK inhibition. In individual comparisons versus undifferentiated controls, SERPINB2 was upregulated during chondrogenesis and downregulated during osteogenesis, confirming the bidirectional lineage switch predicted by loss-of-function experiments [16,17] and growth plate immunohistochemistry [18]. We propose that SERPINB2 functions as a proteolytic brake: by inhibiting uPA and downstream plasmin-mediated MMP activation, SERPINB2 protects the nascent cartilage ECM from premature degradation, stabilising the chondrocyte phenotype. The coordinated upregulation of PLAU (FC +7.63) and PLAUR (FC +2.16) alongside SERPINB2 is consistent with this model—the system simultaneously activates both the protease (uPA) and its inhibitor (SERPINB2), achieving fine-tuned proteolytic control rather than wholesale activation or suppression.

### Context within the OARSI literature

Our findings extend a growing evidence base within the OA biomarker field. Boeuf *et al.* [8] first identified SERPINA1 (356-fold enrichment versus MSCs) and SERPINA3 as chondrogenic markers via a 30,000-gene screen. Our study extends this from clade A to clade B serpins and from static tissue comparisons to the active differentiation process. This convergence is reinforced by recent findings: SERPINF1 as a fibrosis-associated chondrocyte marker driven by EGFR/RhoGTPase signalling [11]; multiple Serpina1 isoforms in a murine chondrocyte-specific transcriptomic signature [24]; serpins among the most reproducibly detected protein families in chondrocyte secretome proteomics [13]; and SERPINE2 ranked among top DEGs in the OARSI single-cell atlas standardisation initiative [10]. The comprehensive review by Paves *et al.* [14] concluded that serpins remain underappreciated in OA despite routine detection in ‘omic analyses. Our systematic cross-platform characterisation directly addresses this gap.

### Signalling context of the SERPIN programme

The coordinated suppression of Wnt antagonists SFRP1 and SFRP4, upregulation of the Wnt effector LEF1, and the PDE4B/PRKACB reconfiguration observed in the Clariom D dataset delineate a multi-pathway signalling context for the SERPIN programme. The suppression of AP-1 components (JUN, FOS, JUNB) during simultaneous SERPINB2 upregulation is mechanistically informative. Elsafadi *et al.* [16] showed that TGF-β1 regulates SERPINB2 through JNK/c-JUN, yet in our KGN-induced model, SERPINB2 is robustly upregulated while JUN and FOS are markedly suppressed. The Bone versus Cart comparison reveals JUN enrichment in osteoblasts relative to chondrocytes (log^2^FC = −1.58, adjusted *P* < 0.05), providing a reciprocal framework: JUN-high/SERPINB2-low defines the osteogenic state, while JUN-low/SERPINB2-high defines the chondrogenic state. This suggests that KGN achieves SERPINB2 upregulation through a mechanism requiring c-JUN suppression—a hypothesis that warrants pathway-specific inhibitor experiments.

### SERPINA9 as a context-dependent marker

The divergent expression of SERPINA9—upregulated in ATCC MSCs but absent in SINREG MSCs—identifies it as a cell-source-dependent marker. This dissociation has practical utility: the SERPINB2/SERPINA9 combination could provide a two-component readout in which SERPINB2 serves as a constitutive chondrogenic marker and SERPINA9 as a transition-phase indicator whose expression depends on cell source characteristics.

### Limitations

Several limitations should be acknowledged. The novel SERPIN candidates (SERPINE2, SERPING1, SERPIND1, SERPINE1) await protein-level validation; Western blot confirmation of SERPINB2 is in progress and will be included prior to journal submission. The *in vitro* 2D monolayer model does not fully recapitulate the three-dimensional joint microenvironment. Clariom D sample size is limited (*n* = 3 per group). Platform differences (∼135,000 vs ∼26,000 transcripts) may contribute to some cross-platform discordances. While converging evidence supports a functional role for SERPINB2, direct loss-of-function experiments in our KGN model have not yet been performed.

### Future directions

Priority experiments include: (i) ELISA quantification of SERPINB2 in OA patient synovial fluid and serum across disease stages; (ii) siRNA knockdown of SERPINB2 in KGN-treated hMSCs to assess effects on chondrogenic markers and the uPA/plasmin cascade; (iii) immunohistochemistry of SERPINB2 in OA versus healthy human cartilage; (iv) longitudinal clinical cohorts correlating circulating SERPINB2 with radiographic OA progression; and (v) three-dimensional chondrogenic models (pellet culture, scaffold-based systems) to confirm findings beyond 2D monolayer.

In conclusion, this multi-platform study validates SERPINB2 as a robust chondrogenic biomarker candidate reproducible across independent cell sources and transcriptomic platforms. The ∼45-fold chondro-osteogenic polarisation, together with converging mechanistic evidence from three independent laboratories, positions SERPINB2 as a *bona fide* lineage-fate determinant. The coordinated remodelling of the Wnt/β-catenin, cAMP/PKA, and uPA/plasmin pathways provides a rich signalling context with direct implications for OA biomarker development and therapeutic targeting of the serpin proteolytic programme.

## DATA AVAILABILITY STATEMENT

The Clariom D microarray dataset has been deposited in the NCBI Gene Expression Omnibus (GEO) under accession number GSE325419. The HGU133+2 dataset is available under accession number GSE325268. Processed differential expression tables are provided as Supplementary Data.

## ACKNOWLEDGMENTS

The authors thank Servicios QSAR, Instituto de Química, Universidad Nacional Autónoma de México (UNAM), for analytical and computational support. The authors gratefully acknowledge SINREG Laboratories (Guadalajara, Jalisco, Mexico) for providing the independent hMSC source and microarray processing used in Dataset 2. We thank Facultad Mexicana de Medicina, Universidad La Salle México, for institutional support.

## AUTHOR CONTRIBUTIONS

B.E.G.R.: Investigation, Resources. E.H.L.: Investigation, Resources. G.L.G.: Investigation. M.C.H.C.: Investigation. A.G.G.R.: Investigation. L.L.P.R.: Investigation. C.E.M.: Investigation. I.R.-B.: Writing – Review & Editing. R.M.V.-G.: Writing – Review & Editing. M.E.E.: Writing – Review & Editing. G.A.M.-N.: Methodology, Writing – Review & Editing. K.M.-M.: Resources, Supervision, Writing – Review & Editing. M.C.-L.: Methodology, Validation, Writing – Review & Editing. J.G.-M.: Conceptualisation, Methodology, Formal Analysis, Investigation, Writing – Original Draft, Visualisation, Supervision, Project Administration, Funding Acquisition.

